# Pathology-defined cell states reveal reproducible transcriptomic signatures across ALS cortical single-nucleus RNA-seq studies

**DOI:** 10.64898/2026.08.07.743523

**Authors:** Charlotte H. van Dijk, Sam Bonsall, Alice Giani, Ryan J. H. West, Jack Humphrey, R. Jeroen Pasterkamp, Johnathan Cooper-Knock, Kevin P. Kenna

## Abstract

Amyotrophic lateral sclerosis (ALS) is a genetically and biologically heterogeneous neurodegenerative disease in which distinct pathogenic mechanisms operate across patients while overt molecular pathology is confined to only a subset of cells. Such features would act to dilute disease-associated transcriptomic signals and complicate the identification of reproducible molecular signatures across the growing number of ALS single-nucleus RNA sequencing (snRNA-seq) studies. Here, we systematically assessed cross-study reproducibility across four cortical ALS snRNA-seq datasets comprising 140 donors (87 ALS) and tested whether pathology-defined cell states improve detection of conserved molecular signatures. Cell-type annotations were harmonized prior to comparison of cell-type-specific pseudobulk differential expression using gene-level, pathway-level, gene-ranking and alternative polyadenylation analyses. We further examined nuclei exhibiting TDP-43 pathology, identified by expression of the *STMN2* cryptic exon. Conventional ALS-versus-control analyses showed limited reproducibility, with minimal overlap of differentially expressed genes or enriched pathways, while fold-change patterns clustered predominantly by study rather than cell type or brain region. Nevertheless, gene-ranking analyses identified reproducible neuronal transcriptional programs, suggesting that biological signal is present but incompletely resolved by current cohort sizes. In contrast, *STMN2* cryptic exon-positive nuclei showed substantially greater concordance, revealing robust TDP-43-associated signatures that partially overlapped independent models of TDP-43 dysfunction while also identifying motor cortex-specific changes, including reduced expression of the recently identified ALS risk gene *UNC13C*. Reproducible ALS-associated alternative polyadenylation changes were not detected, likely reflecting the higher dimensionality and sparsity of polyadenylation site analyses. Together, our findings demonstrate that pathology-defined cell states provide a more reproducible framework for studying ALS transcriptomic alterations than conventional case-control comparisons. We additionally provide an interactive browser to facilitate exploration and comparison of ALS snRNA-seq datasets.

## Introduction

Amyotrophic lateral sclerosis (ALS) is a neurodegenerative disease characterized by progressive motor neuron loss in the motor cortex and spinal cord. Approximately 50% of ALS patients also develop cognitive symptoms, and up to 20% develop frontotemporal dementia (Chiò et al. 2019) Both cell-autonomous and non-cell-autonomous pathogenic mechanisms contribute to motor neuron degeneration in ALS. A pathological hallmark shared by almost all ALS cases is the mislocalization and aggregation of the RNA-binding protein TDP-43 in affected neurons (Mackenzie et al. 2007) Loss of nuclear TDP-43 results in widespread RNA processing defects, including cryptic splicing and alternative polyadenylation, both of which have been implicated in disease pathogenesis (Zeng et al. 2025; Liu et al. 2019; Brown et al. 2022; Newton et al. 2024; Bryce-Smith et al. 2025; Ma et al. 2022) In parallel, increasing evidence supports important contributions of non-cell-autonomous processes to disease progression (Vahsen et al. 2023; Kang et al. 2013; Nagai et al. 2007; Zhang et al. 2026)

Single-nucleus RNA sequencing (snRNA-seq) has become an important approach for dissecting cell-type-specific disease mechanisms directly in patient tissue, leading to the generation of several datasets primarily spanning the motor and frontal cortices of ALS patients (Li et al. 2023; Pineda et al. 2024; Gittings et al. 2023; Bonsall et al. 2026). These studies have provided valuable insights into vulnerable neuronal populations, including deep-layer excitatory neurons such as layer 5 extratelencephalic (L5 ET) neurons, which comprise the upper motor neurons. At the same time, the sparsity inherent to snRNA-seq poses challenges for detecting disease-associated transcriptional changes, particularly when effect sizes are modest or cohort sizes are limited (Squair et al. 2021; Nakatsuka et al. 2025). Consequently, the extent to which reported molecular alterations are reproducible across independent studies remains incompletely understood.

Here, we systematically assess the reproducibility of cell-type-and cell-state-specific transcriptional changes across published ALS cortical snRNA-seq datasets. We combine pseudobulk differential expression analysis, which is less prone to false positives than cell level analyses, with complementary gene-ranking approaches that are less sensitive to outliers and differences in dataset size (Squair et al. 2021; Andreatta and Carmona 2021) We further test whether focusing on nuclei exhibiting a well-established marker of TDP-43 pathology reveals more consistent disease-associated transcriptional programs, and investigate whether alternative polyadenylation changes are reproducibly detected across studies. Finally, we provide our results through an open-access resource to facilitate exploration and comparison of snRNA-seq datasets by the ALS community.

## Results

### Gene and pathway level reproducibility of cell-type-specific transcriptional dysregulation is low

To determine which cell-types are transcriptionally most disrupted in the cortex of ALS patients, we performed differential expression analysis using data from four different snRNA-seq datasets (**FIGURE 1A**). These datasets cover the motor and frontal cortex, totaling 87 ALS donors and 53 controls. To ensure cell-type annotation was consistent, we annotated the nuclei of each dataset using the human cortex reference by the Allen brain institute (**Supplementary figure 1**) (Bakken et al. 2021) A pseudobulk differential expression analysis per cell-type per dataset was performed using DESeq2 (Love et al. 2014) We included all major cell types (astrocytes, oligodendrocytes, oligodendrocyte precursor cells (OPC), microglia, glutamatergic and GABAergic neurons) as well as deep layer neurons and specifically L5 ET neurons as these were previously identified as most disrupted in ALS (Pineda et al. 2024) No one cell-type was consistently the most disrupted across datasets either in terms of number of differentially expressed genes (DEG) or in competitive gene set analyses (**FIGURE 1B, Supplementary table 1-2**). Differences in cohort size across datasets can influence the sensitivity of DEG detection. Given its comparatively small cohort (6 ALS, 6 controls), we excluded the Li et al. dataset from further DEG comparisons.

**Figure 1.**
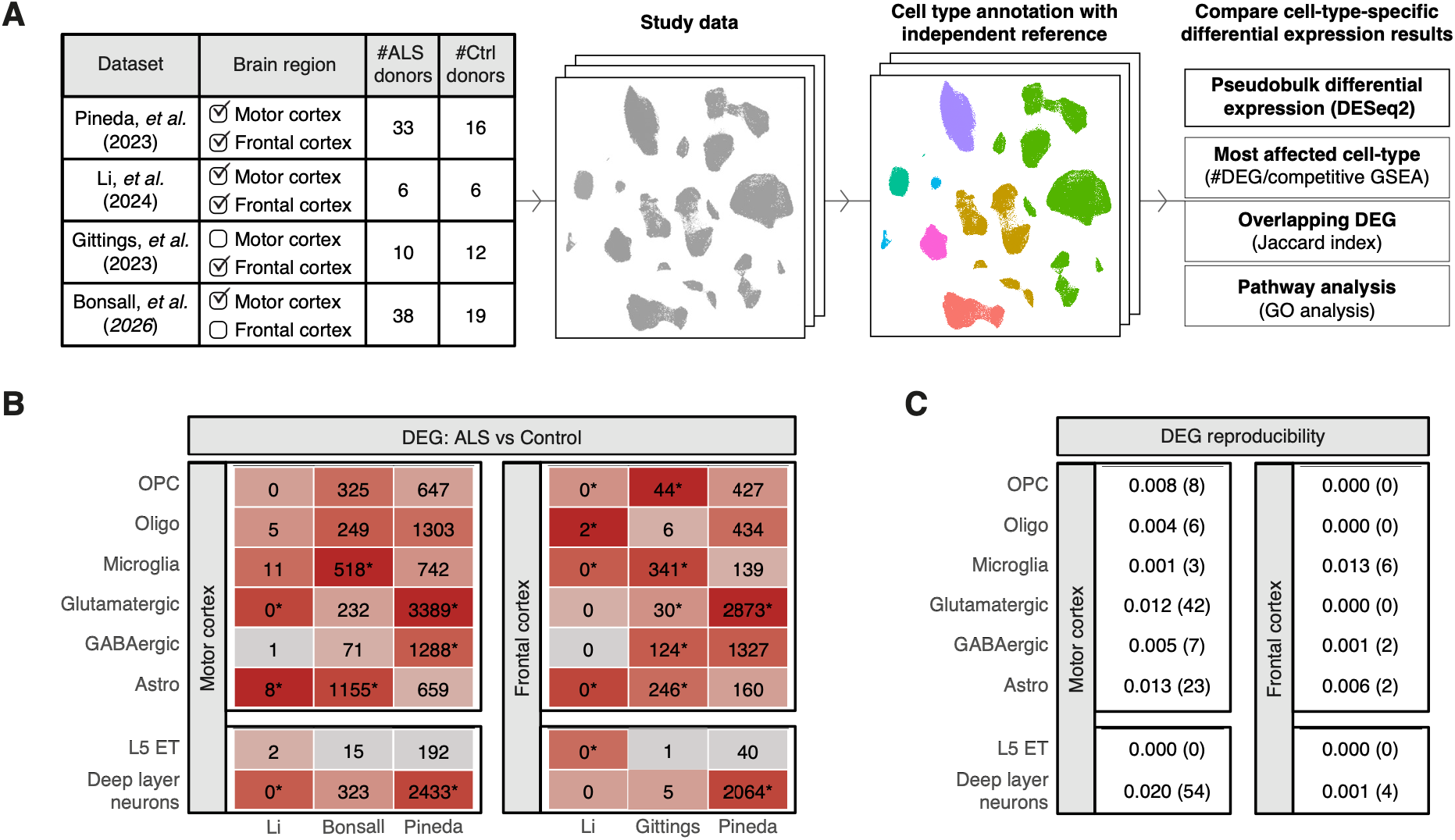
Reproducibility of ALS-induced DEG across cortical snRNA-seq datasets. (**A**) Overview of the datasets included in this study and method for determining reproducibility of cell-type-specific transcriptional changes across datasets. (**B**) Number of cell-type-specific DEG in each dataset derived from a pseudobulk differential gene expression using DEseq2 and colored by enrichment estimates from a competitive gene-set analysis. Boxes with an asterisk indicate cell-types with a significant enrichment (p<0.05) for significant DEG in a competitive gene set analysis. (**C**) Jaccard indices and number of shared DEG in parentheses when comparing cell-type-specific DEG in the motor cortex (comparing Pineda to Bonsall) and frontal cortex (comparing Pineda to Gittings).

Next, we set out to determine which genes are consistently transcriptionally altered in ALS patients. We compared DEG across datasets in a cell-type-and cortical-region-specific manner. To determine global DEG reproducibility, we used the Jaccard index, defined as the number of shared DEG divided by the total number of unique DEG identified in both studies, for each comparison. As a positive control, we confirmed the consistency of cell-type-specific markers in healthy donors across datasets (Jaccard indices ranging between 0.56-0.71, **Supplementary table 3**). However, when comparing disease-associated changes, the reproducibility of individual DEG was extremely low, with Jaccard indices ranging between 0-0.02 (**FIGURE 1C, Supplementary table 4**). To rule out confounders, we repeated the analysis with correction for available donor characteristics (gender, age of the donor, and sequencing batch, **Supplementary table 5**) yet the reproducibility remained low (Jaccard indices ranging between 0.00-0.01).

Finally, we tested whether specific pathways are consistently disrupted in a cell-type-specific manner. We performed Gene Ontology (GO) analysis to identify enriched biological processes among the significant genes from the ALS versus control comparison in each cell type across the two larger motor cortex datasets. In total, out of 414 GO terms that were enriched across the two datasets and six cell types, deep layer and L5 ET neurons, only two GO terms could be reproduced (**Supplementary table 6**).

### Conserved shifts in neuronal transcriptome suggest limited statistical power rather than absence of biological signal hinders reproducibility

To understand why differential gene expression showed limited cross-study reproducibility, we considered two non-mutually exclusive explanations. First, many signals do not reproduce because they reflect study-specific technical or cohort effects. Second, shared transcriptional changes exist but are too subtle for replication across current ALS snRNA-seq studies due to insufficient sample size and a lack of statistical power. To explore these possibilities, we moved beyond canonical gene-level analyses and instead examined the correlation structure of transcriptome-wide case-control changes within and across studies.

If technical or cohort effects dominate the data, it could be expected that observed case-control fold-changes may cluster more by study rather than by biological characteristics such as cell type or brain region. Hierarchical clustering of log-transformed fold changes revealed precisely this pattern, with samples grouping predominantly by study instead of biological characteristics (**FIGURE 2A**). These findings support that even with correction for standard technical and biological confounders during gene level analyses and setting aside questions of power in a second replication dataset, study-specific effects remain an important contributor to observed transcriptional differences.

**Figure 2.**
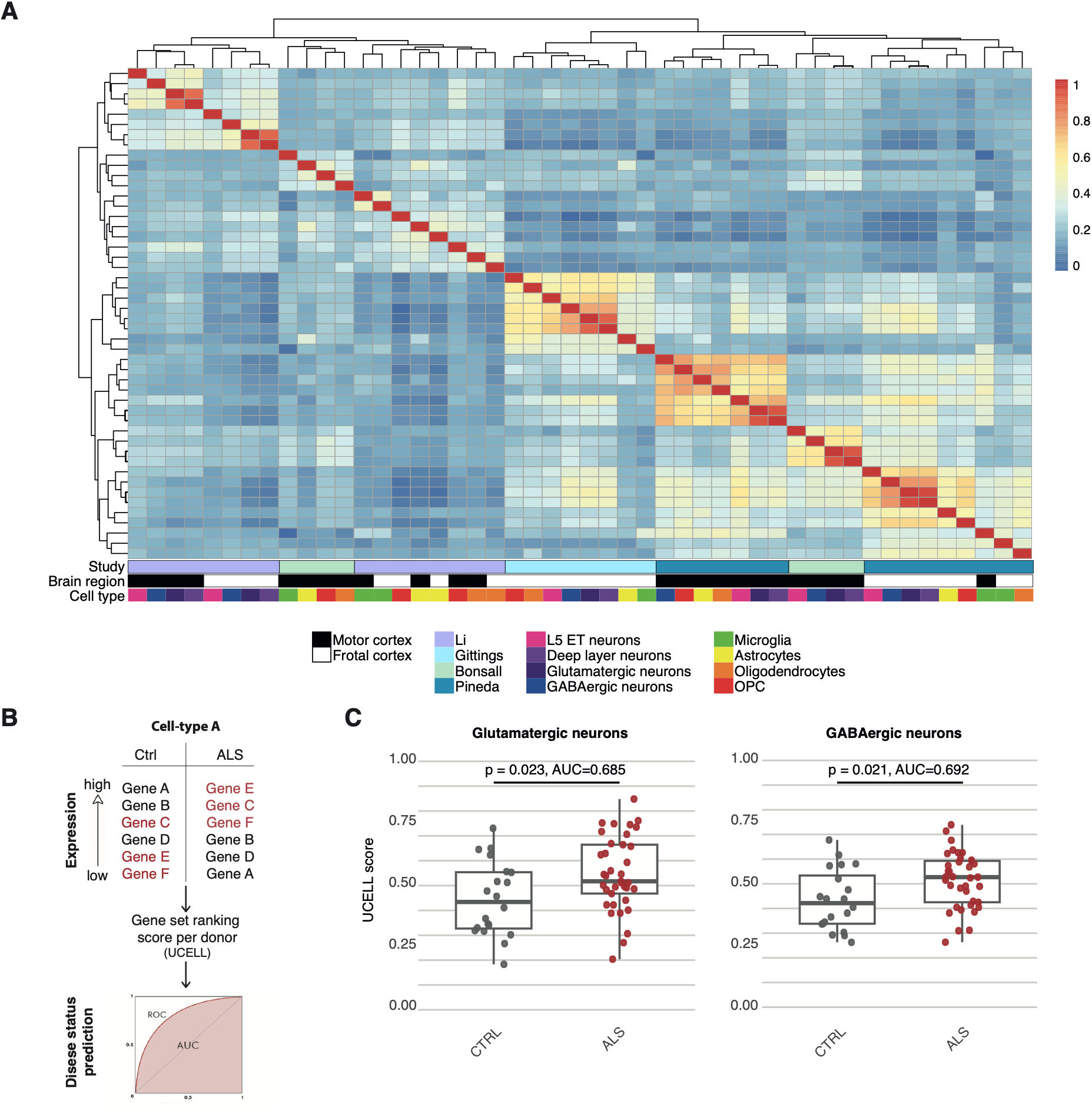
Statistical power as well as dataset specific factors influence reproducibility. (**A**) Correlation matrix heatmap of log2 fold changes derived from cell-type-specific ALS versus control comparison shows clustering based on dataset rather than cell-type or brain region. (**B**) Method for predicting disease status based on the ranking of cell-type-specific ALS-associated DEG (genes in red) and a ranking score is calculated and leveraged to predict case-control status. (C) Glutamatergic and GABAergic UCELL scores derived from the DEG of the Pineda dataset predict case-control status in the Bonsall dataset. P-values are derived from a binomial generalized linear model.

We next evaluated whether limited statistical power, rather than the absence of shared disease-associated transcriptional changes, also contributes to low gene-level reproducibility. To this end, we scored cell type-level pseudobulk samples based on the relative ranking of genes using UCELL (**FIGURE 2B**) (Andreatta and Carmona 2021) Because UCELL summarizes transcriptomic changes across many genes into a single score, it does not provide gene-level resolution but improves sensitivity for detecting coordinated expression shifts across datasets with restricted sample sizes. To assess cross-study reproducibility, we used DEG identified in one dataset (discovery) to generate UCELL scores in an independent dataset (replication). These scores were then used as input for a binomial generalized linear model to predict case–control status, as previously described (Nakatsuka et al. 2025). Model performance was evaluated by calculating area under ROC curves, with the Pineda dataset serving as the discovery cohort and the Bonsall dataset as the independent replication cohort.

As a proof of concept, we validated that UCELL scores can predict disease status of donors in the discovery dataset with ROC-derived AUC values ranging between 0.93 and 1.00 (**Supplementary table 7**). We next investigated whether these gene sets can also be used to predict case-control status in the independent replication dataset. For excitatory (glm p-value = 0.023, AUC = 0.685) and inhibitory neurons (glm p-value = 0.021, AUC = 0.692) the UCELL scores were predictive of disease status (**FIGURE 2C**). For the glia subtypes, no significant relationship could be found (**Supplementary table 8**). These findings indicate that, although individual differentially expressed genes show limited cross-study reproducibility, broader neuronal transcriptional shifts are conserved across independent datasets. Together with the clustering analysis, these results suggest that while study-specific effects are a factor in the limited reproducibility of certain DEG, limited statistical power is also important. Moreover, biologically meaningful neuronal signatures are indeed present and could be further resolved using larger sample sizes.

### Nuclei flagged with TDP-43-associated disease marker show consistent transcriptional disruption

Having established that a conserved neuronal transcriptional signal can be detected across studies, we next asked whether focusing on pathology-defined cell states could further improve the reproducibility of disease-associated gene-level changes. We hypothesized that in nuclei with TDP-43 pathology the transcriptional changes will be severe and therefore might show higher reproducibility of DEG across datasets.

To identify nuclei with TDP-43 pathology, we used expression of the *STMN2* (*chr8:79611214-79616822*, GRCh38) and *KALRN* (*chr3:124701255–124702038*, GRCh38) cryptic exon junctions as a proxy, as previously described (Gittings et al. 2023) We focused our analysis on the motor cortex as this brain region showed the most transcriptional disruption across datasets (**FIGURE 1B**). *STMN2* cryptic junction reads were exclusively found in ALS samples and absent from controls in all three datasets (**FIGURE 3A, Supplementary figure 2A**). Although *KALRN* cryptic junction reads were identified, they were also present in controls with no significant increase in ALS donors (P=0.31 (Pineda), P=0.49 (Bonsall), P=0.39 (Li), Wilcoxon test) (**FIGURE 3A, Supplementary figure 2A**). We therefore focused our analyses on the *STMN2* cryptic junction, with the vast majority of *STMN2* cryptic exon junction positive (STMN2CE^+^) nuclei originating from glutamatergic neurons and in particular the L2/3 intratelencephalic (IT) subclass (**Supplementary figure 2B**).

**Figure 3.**
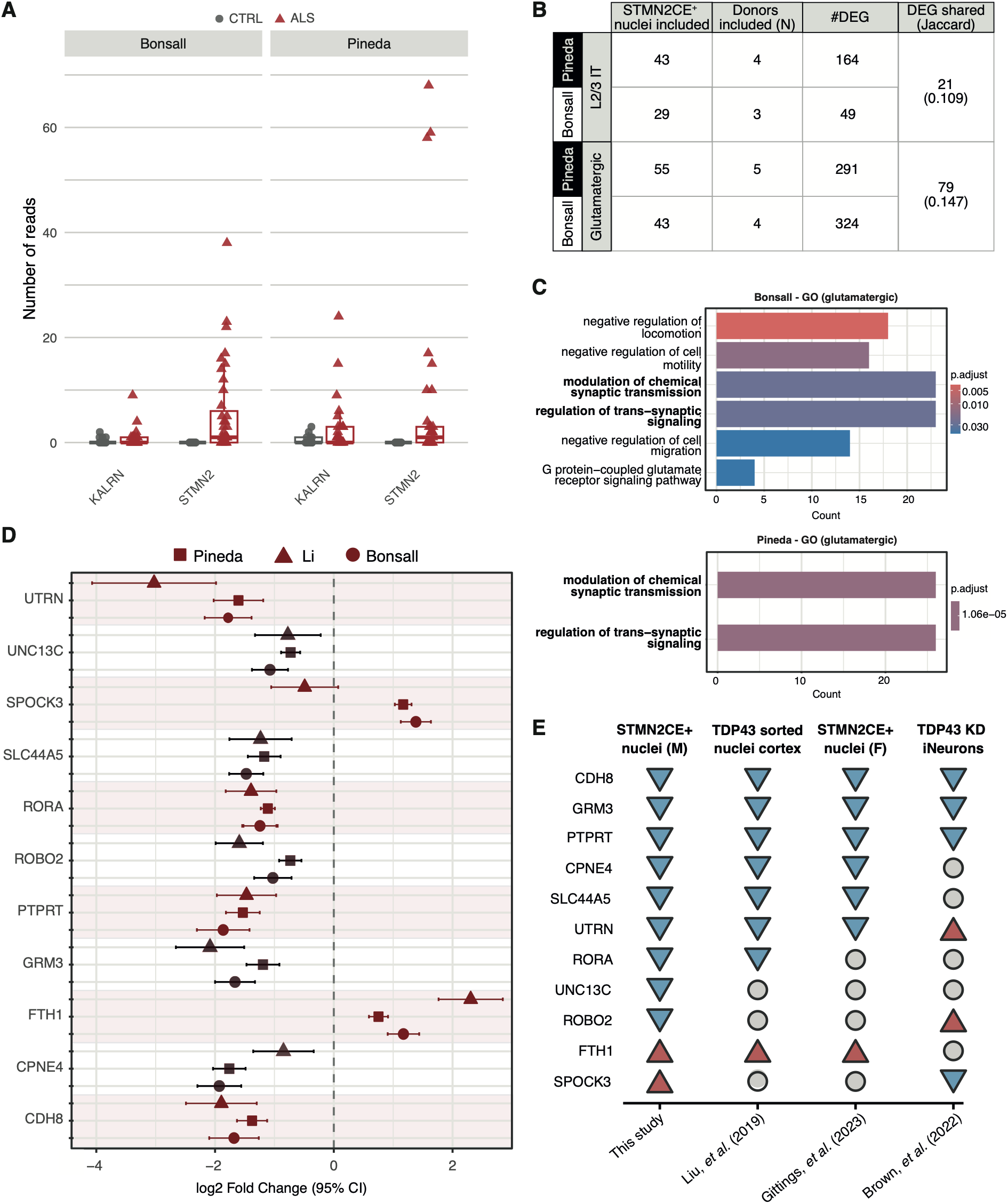
Differential gene expression analysis comparing STMN2CE+ to STMN2CE-neurons shows consistent dysregulation across datasets. (**A**) Number of cryptic junction-spanning reads identified in the motor cortex data of each donor of the Bonsall and Pineda dataset. The cryptic junctions are defined as chr8:7SC11214-7SC1C822 for STMN2 and for KALRN chr3:124701255–124702038 (GRCh38). (**B**) Differential gene expression analysis for STMN2CE+ vs STMN2CE-glutamatergic or specifically L2/3 IT neurons. Depicted are the number of donors and STMN2CE+ nuclei included in each analysis, the number of DEG, and the number of DEG shared with the Jaccard indices in brackets when comparing the results between the Bonsall and Pineda dataset. (**C**) GO terms derived from a gene ontology (biological process) gene set enrichment analysis of DEG identified by comparing glutamatergic STMN2CE+ vs STMN2CE-nuclei for both the Bonsall dataset (top) and Pineda dataset (bottom). (**D**) Forest plot representing the log2fold changes of the eleven genes shared by the glutamatergic and L2/3 IT STMN2CE+ vs STMN2CE-comparison in both the Bonsall and Pineda dataset. As the Li dataset had only two donors with sufficient STMN2CE+ glutamatergic nuclei, the differential expression results were only used to compare direction of effect. (**E**) Comparison of the eleven genes to differential gene expression in other TDP-43 pathology models. The blue inverted rectangles represent significant downregulation, the red upward facing rectangles represent significant upregulation and the grey circles indicate no significant change.

To determine the reproducibility of transcriptional changes in nuclei with TDP-43 pathology, we performed pseudobulk differential expression analysis comparing STMN2CE^+^ and STMN2CE^-^ nuclei across all glutamatergic neurons as well as the L2/3 IT subclass. Despite the sparsity of STMN2CE⁺ nuclei, Jaccard indices across datasets were five to seven times higher than the highest cell-type level comparison (0.11 and 0.15 for STMN2CE^+^ L2/3 IT and STMN2CE^+^ glutamatergic neurons, respectively) (**FIGURE 3B**). GO analysis of the DEG from STMN2CE^+^ glutamatergic neuron analysis also showed two overlapping pathways out of the six possible overlapping terms (**FIGURE 3C**). These findings confirm that focusing the differential expression on nuclei with a disease marker increases the reproducibility of DEG across datasets.

Eleven DEG, nine downregulated and two upregulated, were shared between the L2/3 IT and glutamatergic neuron STMN2CE^+^ versus STMN2CE^-^ comparison. The Li dataset did not include sufficient STMN2CE^+^ nuclei to support a robust differential expression analysis. Nevertheless, we used the two donors that had sufficient STMN2CE^+^ glutamatergic nuclei to examine concordance in up-and downregulation. Out of the eleven overlapping genes, ten showed the same direction of effect in the Li dataset (**FIGURE 3D**). *PTPRT* is a known splice target of TDP-43 (Ma et al. 2022) and *FTH1* has previously already been shown to be transported by TDP-43 to axons where it regulates iron homeostasis and oxidative stress (Jinno et al. 2025)

Recently, it was shown that the effect of TDP-43 pathology on RNA expression and processing is tissue-and model-system-specific(Newton et al. 2024) Therefore, we wondered whether the eleven genes that showed consistent dysregulation would also reproduce across other TDP-43 pathology model systems. We compared the results to three other datasets: STMN2CE^+^ versus STMN2CE^-^ nuclei from the frontal cortex, TDP-43-sorted nuclei from the postmortem cortex of ALS and FTD donors, and a TDP-43 knockdown model in iNeurons (**FIGURE 3E**) (Liu et al. 2019; Brown et al. 2022; Gittings et al. 2023) Overall, *CDH8, GRM3,* and *PTPRT* showed consistent downregulation across all TDP-43 models. Additionally, *CPNE4*, *SLC44A5*, *UTRN* and *FTH1* showed consistent dysregulation in postmortem brain tissue. The iNeuron model where TDP-43 is knocked down showed the least overlap, whereby some genes (*UTRN, ROBO2, SPOCK3*) even showed dysregulation in the opposing direction. *RORA*, *UNC13C*, *ROBO2*, and *SPOCK3* dysregulation was specific to the motor cortex. Interestingly, *UNC13C* was recently discovered as a putative ALS risk gene (Hop et al. 2026)

### Larger datasets are required for detection of reproducible polyadenylation site switching

Recently, cryptic alternative polyadenylation (APA) has been identified as a potential disease mechanism in ALS (Bryce-Smith et al. 2025; Zeng et al. 2025) Cell-type-specific APA changes were also identified at a single-cell level in the orbitofrontal cortex of ALS patients (McKeever et al. 2025). We therefore examined whether cell-type-specific APA disruptions could be reproducibly detected in the ALS motor cortex. Because the 3′ Chromium chemistry used across these datasets generates reads that pile up close to the polyadenylation site, we used the splice-aware peak-calling tool Sierra to identify polyadenylation sites (PAS) (**Supplementary figure 3A**), with the majority of sites mapping to the 3′ UTR consistent with known PAS distributions (**Supplementary figure 3B**) (Patrick et al. 2020) PAS usage was reproducible across cell types and cell classes in healthy donors, validating our approach (**Supplementary figure 3C**). However, the number of differentially expressed PAS between ALS and control varied markedly by dataset (**Supplementary figure 3D**). UCELL scores derived from Pineda’s differential PAS successfully predicted case-control status within the discovery dataset (**Supplementary figure 3E**) but had no predictive power in the independent Bonsall dataset (**Supplementary figure 3F**). Given that pathology-defined cell states improved the reproducibility of gene-level changes, we tested whether this also held for APA. Comparing STMN2CE^+^ versus STMN2CE^-^ glutamatergic neurons in the Pineda dataset identified four significant PAS (padj < 0.1) which cumulatively involved 3′ UTR shortening in *RPS6KA2* and 3′ UTR lengthening in *CADM3* (**Supplementary table 9**). *CADM3* is biologically interesting, having previously been proposed as a genetic risk factor for Charcot Marie Tooth disease type 2 and a candidate ALS biomarker in CSF (Rebelo et al. 2021; Muqaku et al. 2025). However, no significant APA changes were detected in the Bonsall dataset (**Supplementary table 10**). Together, these findings suggest that while our approach can nominate promising candidate APA targets, larger cohorts will likely be needed to establish robust cell-type-and cell-state-specific APA changes in ALS.

### An interactive resource to facilitate cross-study exploration of cell-type-specific gene expression changes in ALS

To facilitate exploration of the results generated in this study, we developed an interactive browser for the ALS research community (https://chvandijk.shinyapps.io/ALS_snRNA_Reproducibility). The browser enables users to query genes and directly compare cell-type and cell-state specific differential expression across the four cortical snRNA-seq datasets analyzed here. Effect sizes, statistical significance, and cross-study concordance can be viewed side-by-side, allowing rapid assessment of whether transcriptional changes are consistently observed or study specific.

## Discussion

This study systematically evaluated the reproducibility of cell-type-and cell-state-specific transcriptional changes across ALS cortical snRNA-seq datasets. Our analyses show that pathology-defined cell states, identified through expression of the *STMN2* cryptic exon, reveal robust and reproducible transcriptional signatures across studies. In contrast, conventional case-control differential expression yielded limited overlap at the individual gene level. Gene-ranking analyses further revealed reproducible signatures in neurons, indicating that disease-associated signal is present in these datasets but will require larger cohorts to be reliably resolved. Together, these findings provide both a framework for interpreting existing ALS snRNA-seq studies and a roadmap for improving future analyses. In addition, we provide an interactive resource to facilitate exploration and comparison of ALS cortical snRNA-seq datasets by the community.

Our observations are consistent with recent benchmarking studies in other neurological disorders, including Alzheimer’s disease and schizophrenia, where cross-study reproducibility of single-cell differential expression has likewise proven challenging despite the availability of increasingly large datasets (Nakatsuka et al. 2025; Mathys et al. 2023) In Alzheimer’s disease, reproducible transcriptional signatures became more apparent as cohort sizes expanded into the hundreds of donors. Our gene-ranking analyses suggest that ALS transcriptional effect sizes are of a similar magnitude, indicating that larger cohorts are likely needed to improve the detection of reproducible cell-type-specific changes. This challenge is likely to be even greater for alternative polyadenylation, where the higher dimensionality of PAS-based analyses further reduces statistical power. Although we were unable to identify reproducible APA changes across current datasets, our analyses nominate *CAMD3* as a candidate event for further investigation. Together with the growing evidence implicating cryptic APA in ALS pathogenesis, our investigation thus calls for follow-up analyses of APA in larger future cohorts.

Our results further demonstrate that defining biologically meaningful disease states represents a compelling strategy for increasing reproducibility without necessarily requiring dramatically larger cohorts. Similar approaches have proven successful in other neurodegenerative diseases, such as Huntington’s disease, where transcriptional alterations are primarily observed in neurons carrying somatically expanded CAG repeats, while neurons without expansion show profiles that more closely resemble controls (Handsaker et al. 2025). Increasing cohort sizes to the scale of studies in Alzheimer’s disease will involve substantial challenges, with the availability of sufficient high quality sample material being paramount. Therefore, future studies should additionally focus on developing new strategies to enriching disease-relevant cell states or vulnerable populations to reveal molecular changes that may otherwise be obscured when analyzing heterogeneous patient-derived tissue. For cell types or states lacking clear nuclear markers, including upper motor neurons and reactive glial subtypes, spatial transcriptomic approaches may provide an alternative strategy by preserving the cellular context of disease-associated changes. Such approaches may also help elucidate interactions between vulnerable neurons and their surrounding environment, for example the reported spatial association between reactive microglia and motor neurons in ALS (Jara et al. 2017).

Beyond defining disease-relevant cell states, our findings highlight the importance of studying transcriptional changes in an appropriate tissue and disease context. Several genes, such as *PTPRT,* dysregulated in STMN2CE^+^ nuclei from the motor cortex showed consistent downregulation across independent TDP-43 model systems. Some genes, however, showed dysregulation in the opposite direction in an iNeuron TDP-43 knockdown model, while others were detected exclusively in the motor cortex, showing no significant change in other TDP-43 models, in line with previous work showing that TDP-43 effects are context-dependent (Newton et al. 2024) Among the group of motor cortex-specific genes, *UNC13C* is particularly notable, as it was recently identified as a putative ALS risk gene, lending independent support to the biological relevance of this motor cortex-specific signal (Hop et al. 2026) Our work therefore extends prior knowledge on TDP-43 pathology by identifying transcriptional changes that occur in the patient motor cortex.

A remaining open question is what underlies observed dataset-specific effects. We included available covariates in our analysis to account for standard technical and biological confounders but, yet this did not lead to improved reproducibility. Another possibility is that differences in patient composition between studies contribute to the observed variability. ALS encompasses a broad spectrum of clinical trajectories and genetic etiologies, which may give rise to distinct disease-associated transcriptional changes (Eshima et al. 2023; Swinnen and Robberecht 2014) Differences in donor ancestry could also contribute, as previous studies have shown that genetic background contributes substantially to variation in gene expression between individuals (Oelen et al. 2022; Tung et al. 2017; Kim-Hellmuth et al. 2020) Finally, technical factors such as site of dissection or tissue handling may also contribute, though these remain difficult to assess retrospectively. Identifying the specific sources of dataset-specific artifacts will be important for informing future study design and cross-dataset integration.

In conclusion, this study provides a systematic evaluation of reproducibility across ALS cortical snRNA-seq datasets and identifies pathology-defined cell states as a robust framework for studying disease-associated transcriptional changes. By distinguishing biological signals that are consistently detected across independent studies from those that remain sensitive to study-specific variation, our work provides practical guidance for future experimental design and data interpretation. Notably, our gene-ranking analyses reveal that reproducible disease-associated signal is already present in neurons, indicating that larger cohorts should eventually make this signal detectable at the individual gene level as well. Together with the accompanying interactive browser, these results establish a community resource for exploring and comparing ALS snRNA datasets.

## Materials and methods

### snRNA-seq datasets

For the Pineda dataset, raw sequencing data was obtained from the Sequencing Read Archive (SRA) (BioProject accession PRJNA1073234) and processed data, including count matrices and cell-type-annotations were downloaded from Synapse (Project SynID: syn51105515). For the Li dataset, raw sequencing data was obtained from SRA (BioProject accession PRJNA907949) and processed data from the following Zenodo repository https://doi.org/10.5281/zenodo.8190317. For the Gittings dataset, processed data was downloaded from CELLxGENE (https://cellxgene.cziscience.com/collections/aee9c366-f2fb-470b-8937-577d5d87d3fc). Raw RNA-seq data for the Bonsall dataset will be made available via the National Center for Biotechnology Information’s Gene Expression Omnibus database (Bioproject ID is PRJNA1468995) as described Bonsall et al(Bonsall et al. 2026)

### Cell type reference mapping

The cells were annotated using s a third independent dataset, the Allen brain Human M1 10x, as a reference. The processed count matrix and cell type annotations were downloaded from the data portal (https://portal.brain-map.org/atlases-and-data/rnaseq/human-m1-10x). Since non-neuronal subclasses are shared across brain regions and cortical layer markers are conserved across cortical areas, this reference could be used for both the motor and frontal cortex (Jorstad et al. 2023). This approach enabled direct comparison of cell types between two datasets. The functions FindTransferAnchors and TransferData from the Seurat (v4.4) R package were used to map the cell classes and subclasses.

### Differential gene expression

Differential gene expression was performed using DESeq2 (Love et al. 2014). Counts were aggregated by cell-type or cell-class and donor. Genes with less than 10 counts were excluded. Genes with padj<0.05 were considered significant. Potential confounders were added as covariates to the design formula. Analyses were performed to control for the reported sex of the donors or for any potential confounder in the published sample metadata of the study. For the Pineda study these additional confounders include Braak stage, Thal phase, and age of death. For Li, these include the age at death and for the Bonsall dataset age of death and sequencing batch. Competitive gene set enrichment was performed using the p-values of genes with a basemean>1 of the cell-type-specific DE analysis and compared the p-values of one cell-type to all others using the *geneSetAssoc* function with test = “lm” from the RVAT package (https://github.com/KennaLab/rvat).

### GO analysis

Pathway enrichment analysis was performed using the clusterProfiler (v4.18.2) R package (Yu et al. 2012). The *enrichGO* functions was used with parameters *pvalueCutoff* = 0.05, *qvalueCutoff* = 0.2, *ont* = “BP”, and *pAdjustMethod* = “BH”. As background, genes with a basemean>5 were used.

### Gene ranking and case-control status prediction

To calculate gene ranking scores, the R package UCELL (v2.6.2) was used which applies a Mann-Whitney U statistic to gene set enrichment scores for predefined gene sets (Andreatta and Carmona 2021) These gene sets were defined as the significant genes from cell-type-specific differential gene expression analyses of a discovery dataset (Pineda et al). Within an independent replication dataset (Bonsall et al), per sample enrichment scores were then calculated and used to predict disease labels using a binomial generalized linear model using the *glm* function from the stats (v4.5.1) R package. ROC curves were determined using the *roc* function and the AUC was calculated using the *auc* function, both from the pROC (v1.19.0) R package.

### STMN2CE detection

The method to detect *STMN2* and *KALRN* cryptic junction expression nuclei was based on the publication by Gittings, *et al*. To detect *STMN2* and *KALRN* cryptic junction expressing nuclei, raw fastq files were aligned using the cellranger (v7.1.0) count command with GRCh38-2020-A as reference. Cell barcodes of junction-spanning reads were extracted from the aligned sequencing files using a custom python script that analyses the CIGAR strings of the aligned sequencing reads using the pysam package and identifies reads matching the specified intron coordinates and requiring a minimum contiguous overhang of six nucleotides on both splice junction anchors. The junction locations are *chr8:7SC11214-7SC1C822* and *chr3:124701255–124702038* (GRCh38), for *STMN2* and *KALRN*, respectively.

### STMN2CE DEG analysis

Gene level counts within STMN2CE^+^ nuclei were generated by first selecting reads from STMN2CE^+^ nuclei using the function *filterbarcodes* from sinto (v0.9.0). The reads were then transformed back to fastq files using cellranger’s *bamtofastq* command after which count matrices were generated using cellranger’s *count* as described above. Comparisons of STMN2CE^+^ and STMN2CE^-^ nuclei were performed for only “glutamatergic” and “L2/3 IT” neurons. Additional cell classes were not considered due to limited observations of STMN2CE^+^ nuclei. Our analyses were restricted to donors carrying a minimum of 3 STMN2CE^+^ nuclei. For glutamatergic neurons, nuclei were required to meet all quality control criteria specified in the source studies. For rarer L2/3 IT STMN2CE^+^ neurons we performed additional analyses where the minimal quality control criteria was adjusted to <10 percent mitochondrial reads and>500 expressed features. Gene expression counts were then aggregated in a pseudobulk manner and differential expression analyses performed using DESeq2 and the following design: *∼ Donor + STMN2_status*. Genes with tested for reproducibility using an adjusted p-value threshold of <0.1.

To evaluate reproducibility of DEG derived from our analyses of STMN2CE^+^ versus STMN2CE^-^ nuclei we further compared the results to other models of TDP-43 pathology. DEG derived from STMN2CE^+^ versus STMN2CE^-^ nuclei in the frontal cortex were retrieved directly from the study by Gittings, *et al*. (Gittings et al. 2023). RNA-seq count data from TDP-43 FACS-sorted cortical nuclei of ALS/FTD patients as well as from iNeurons with or without TDP-43 knockdown were used as input for a differential expression analysis performed using DESeq2 (Liu et al. 2019; Brown et al. 2022) Only genes with more than 10 counts were included. Genes were considered significant with an adjusted p-value <0.05.

### PAS identification

Raw sequencing data was aligned to build GRCh38 of the human genome (refdata-gex-GRCh38-2020-A) using Cell Ranger (v7.1.0). Analyses were restricted to nuclei passing the quality control criteria outlined in the source publication, and to reads with a mapping quality >30 which were extracted using sinto (v 0.9.0, https://github.com/timoast/sinto) and SAMtools (v1.18) (Danecek et al. 2021) Internal priming artefacts were removed using polyAfilter (v3) (https://github.com/MarekSvob/polyAfilter), which removes reads mapping 300 bp upstream of a stretch of eleven adenosines, allowing for 2 mismatches. pA site identification was performed using the splice-aware peak-calling tool Sierra (v0.99.27) (Patrick et al. 2020) Read junctions, required as input by Sierra, were called using RegTools (v0.5.2) (Cotto et al. 2023) Peak calling was performed on a per-sample basis using the GENCODE v32 GTF annotation of the human transcriptome, with annotated 3′ UTRs extended by 500 bp to enable detection of distal polyadenylation sites beyond the annotated transcript ends. Sample-level peaks were merged using Sierra’s MergePeakCoordinates function. Only peaks that were found in more than one sample were kept and nested peaks on the same exon were removed using a custom script. PAS counts were obtained using Sierra’s CountPeaks function. PAS that are expressed in less than 30 cells were removed. Location of the PAS was determined using Sierra’s *AnnotatePeaksFromGTF* function.

### APA analysis

APA analyses were restricted to protein-coding genes with 2-10 PAS. PAS counts were aggregated across cell-type and donor in a pseudobulking manner. Analyses were restricted to PAS expressed in 10% of samples. PAS that localized to annotated 5’ UTR were excluded. The functions *estimateSizeFactors*, *estimateDispersions*, *testForDEU*, *estimateExonFoldChanges* were used to test for significant differences in APA across groups. DEXSeq analysis was performed using genes as the groupID and ∼ sample + exon + condition:exon as the design formula whereby exon is a placeholder name referring to PAS. Only PAS with an adjusted p-value below 0.05 were considered significant. For the healthy cell-type comparison, the data was restricted to only non-neurological control donors and condition refers to the cell type that was compared. For the ALS-driven APA analysis, condition refers to disease status. For the STMN2CE^+^ versus STMN2CE^-^ analysis, condition refers to STMN2CE status.

In order to compare results across datasets, the PAS sites were mapped to cleavage sites. The cleavage site of a pA site was defined by overlapping the region 100 bp upstream of the peak termination site with the cleavage sites defined in the PolyAsite3.0 database (Moon et al. 2025) If multiple cleavage sites overlapped, the one with the highest stringency score was selected. For both datasets, over 95% of pA sites were matched to a cleavage site and these were used for subsequent ranking analysis. UCELL scores and case-control prediction were performed as described in the section “*Gene ranking and case-control status prediction*”, with the exception that cleavage sites were used in place of genes.

## Statistical analysis

Statistical analyses were performed using R (R-4.3.1).

## Data access

This study did not generate new sequencing data. All datasets analyzed were previously published; accession numbers and repository details for each dataset are provided in the Methods. The results of our reproducibility analyses, including all cell-type-, dataset-, and gene-level comparisons, can be explored via our browser (https://chvandijk.shinyapps.io/ALS_snRNA_Reproducibility).

## Acknowledgements

K.K. is supported by grants from the Dutch Research Council (grant no. ZonMW-VIDI 91719350) and the ALS Foundation Netherlands. SB, JCK and RW are supported by Tambourine’s ALS Breakthrough Research Fund in partnership with the Milken Institute.

## Authors’ contributions

Study conception and design: KK, CD. Supervision: KK, JP. Data analysis: CD. Data interpretation: CD, KK, JP, SB, RW, JCK, AG, JH. Resources: SB, RW, JCK. Writing – original draft: CD, KK. Writing – review & editing: All authors. All authors read and approved the final manuscript.

## Supplementary material table legend

**Supplementary material table 1 Cell-type-specific DEG across brain regions and datasets.** This table summarizes the significant (padj<0.05) DEG per cell-type per dataset and the direction of effect (up-or downregulated in comparison to control).

**Supplementary material table 2 Competitive gene set enrichment results.** To test which cell-type was most affected in each dataset, we performed a competitive gene set enrichment analysis. As input, the p-values of genes (basemean > 1) derived from the cell-type-specific ALS versus control differential expression analysis were used. The p-values of one cell-type were compared to all others in an iterative manner. Depicted are the estimated regression coefficient (“effect”) and the p-value results of the GSEA.

**Supplementary material table 3 Cell-type DEG comparison in healthy donors.** To assess reproducibility of the cell type identity across datasets, we performed a differential expression analysis comparing one cell type to the rest (“Cell-type versus other comparison”) or else comparing specific cell type pairings (“Cell-type versus cell-type comparison”). Analyses were restricted to nuclei from the motor cortex of healthy donors in the Pineda and Bonsall datasets. The Pineda and Bonsall datasets were analyzed separately and the overlap of DEG between the two summarized as the indicated Jaccard index values.

**Supplementary material table 4 Overlapping DEG between datasets.** This table summarizes the significant DEG (padj<0.05) shared between each dataset and the Pineda dataset and their direction of effect (up-or downregulated in comparison to control).

**Supplementary material table 5 Covariate-controlled DEG comparison**. The cell-type-specific differential analysis and DEG comparison was repeated, this time controlling for the sex of the donor (top) or any potential confounder (bottom, covariates included are depicted in the design formula). The numbers in each study column represent the number of DEG identified using the covariate-controlled analysis.

**Supplementary material table 6 GO term comparison.** The number of enriched GO terms (biological process) amongst the cell-type-specific DEG in the Pineda (motor cortex) and Bonsall datasets, and the number shared by both datasets.

**Supplementary material table 7 UCELL case-control prediction in the Pineda dataset.** Pineda’s cell-type-specific DEG were used to generate UCELL scores in the Pineda dataset that were then used to predict case-control status within the same dataset using a binomial generalized linear model. Depicted are the estimate and the p-value from this prediction and the AUC values.

**Supplementary material table 8 UCELL case-control prediction in the glial cells of the Bonsall dataset.** Pineda’s cell-type-specific DEG were used to generate UCELL scores in the Bonsall dataset that were then used to predict case-control status using a binomial generalized linear model. Depicted are the estimate and the p-value from this prediction and the AUC values.

**Supplementary material table 9 APA analysis results for STMN2CE^+^ versus STMN2CE^-^ glutamatergic nuclei in the Pineda dataset.** The PeakID is defined as follows: gene:start peak-end peak:strand, with 1 being the sense strand and-1 the antisense. The genomic_feature(s) column represents the peak location in the reference genome annotation file.

**Supplementary material table 10 APA analysis results for STMN2CE^+^ versus STMN2CE^=^ glutamatergic nuclei in the Bonsall dataset.** The PeakID is defined as follows: gene:start peak-end peak:strand, with 1 being the sense strand and-1 the antisense. The genomic_feature(s) column represents the peak location in the reference genome annotation file.

## Supplementary figures

**Supplementary Figure 1.**
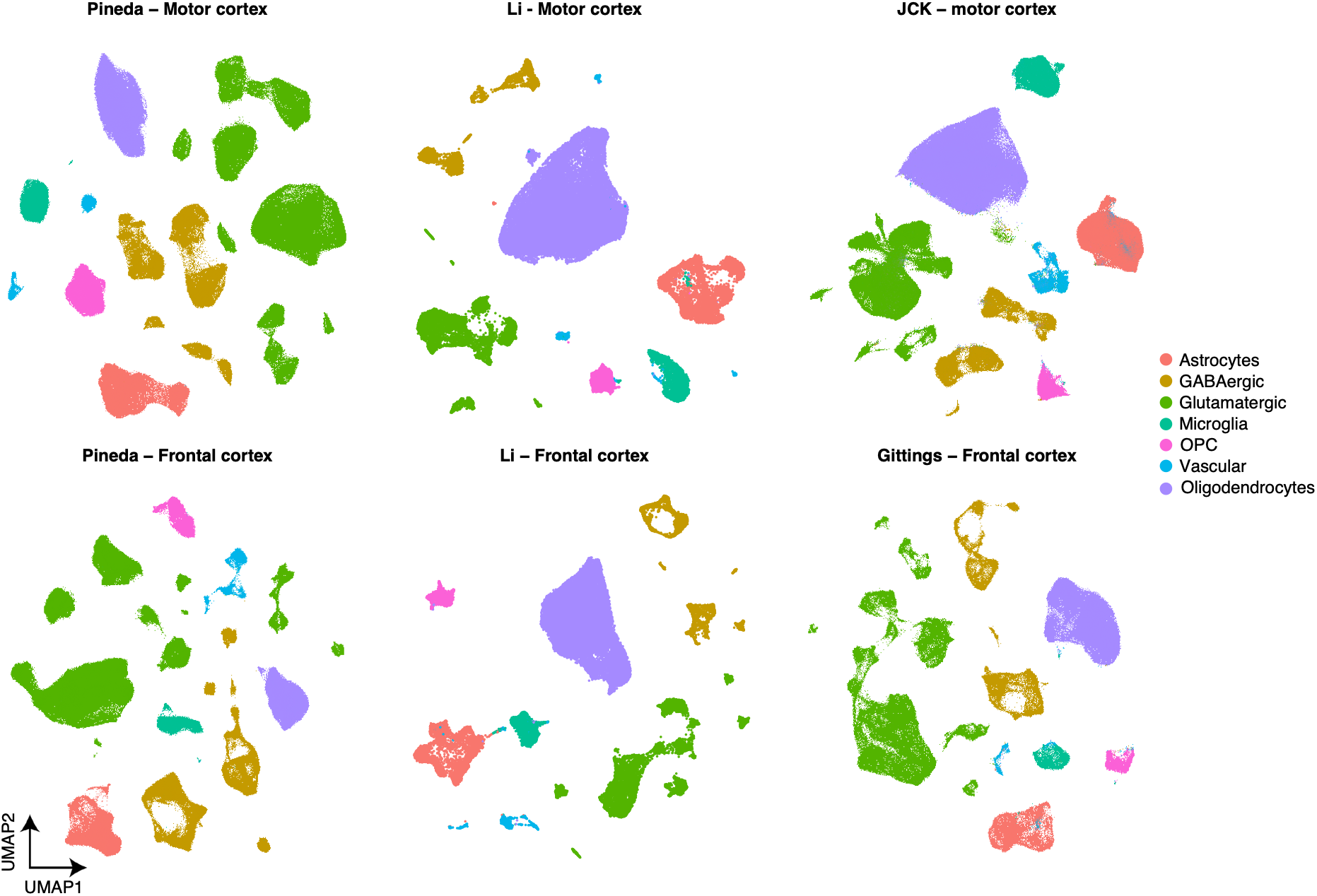
Cell type annotation. UMAP representation of nuclei from each dataset colored by cell type annotation. The Allen Brain cortex atlas was used as a reference.

**Supplementary Figure 2.**
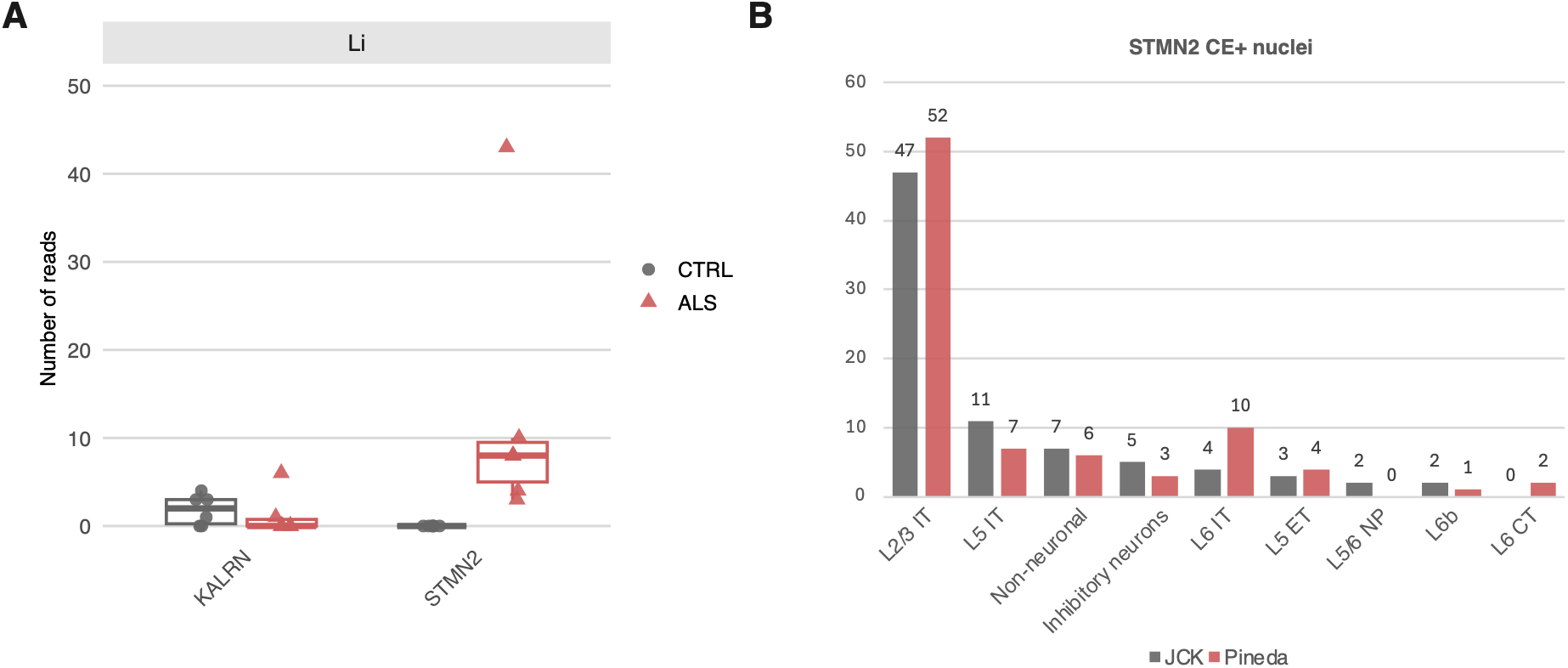
STMN2 cryptic exon. (**A**) Number of cryptic junction-spanning reads identified in the motor cortex data of each donor across all cell-types within the Li dataset. The cryptic junctions are defined as chr8:7SC11214-7SC1C822 for STMN2 and for KALRN chr3:124701255–124702038 (GRCh38). (**B**) Cell type annotation of STMN2CE+ nuclei in the Bonsall and Pineda datasets.

**Supplementary Figure 3.**
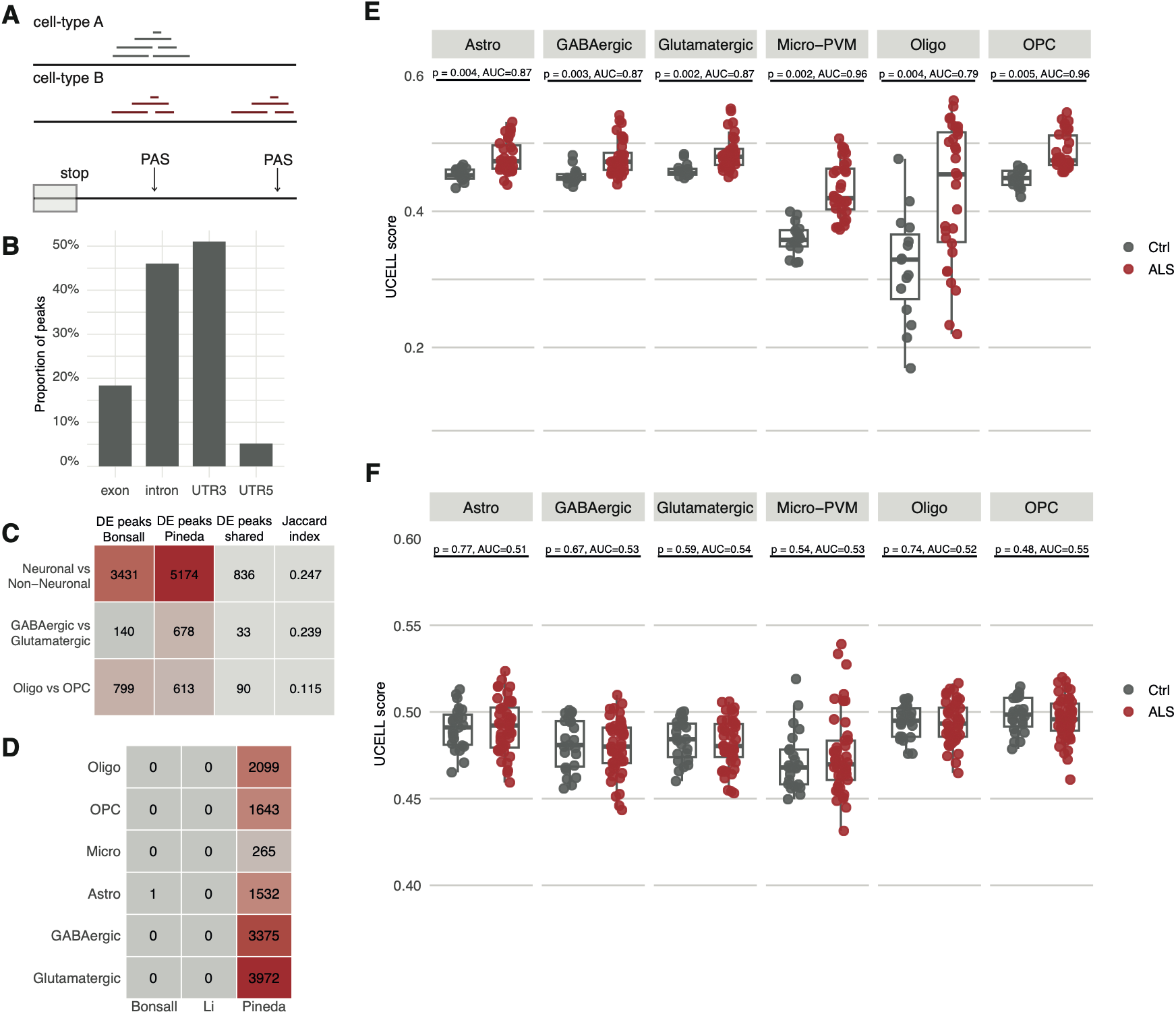
Polyadenylation changes in the motor cortex of ALS patients. (**A**) Schematic representation of the method whereby reads form peaks close to the polyadenylation site (PAS) allowing and by means of a peak-detection tool PAS can be identified. (**B**) Location of identified PAS sites in the Pineda dataset. (**C**) Sensitivity analyses evaluating the number of peaks exhibiting differential expression (DE peaks) across major cell type annotations in healthy control donors from the Bonsall and Pineda datasets. (**D**) Number of DE peaks in the Bonsall, Li, and Pineda dataset when comparing ALS to controls in a cell-type-specific manner. (**E**) UCELL scores derived from the DE peaks of the Pineda dataset predict case-control status across all major call types within the discovery dataset. (**F**) UCELL scores derived from the DE peaks of the Pineda dataset fail to predict case-control status in the Bonsall dataset, indicating a lack of reproducible biological signal across all major cell types. P-values are derived from a binomial generalized linear model.

